# Longitudinal multi-omics along gingivitis development reveal a suboptimal-health gum state with periodontitis-like microbiome

**DOI:** 10.1101/2020.09.26.315127

**Authors:** Shi Huang, Tao He, Feng Yue, Victor Xu, Spring Wang, Pengfei Zhu, Fei Teng, Zheng Sun, Xiaohui Liu, Gongchao Jing, Xiaoquan Su, Lijian Jin, Jiquan Liu, Jian Xu

## Abstract

Most adults experience episodes of gingivitis, which can progress to the irreversible, chronic state of periodontitis. However the mechanistic roles of plaque in gingivitis onset and progression to periodontitis remain elusive. Here, we integrated the longitudinal multi-omics data from plaque metagenome, metabolome and salivary cytokines in 40 adults who transit from naturally-occurring gingivitis (NG), to healthy gingivae (baseline) and then to experimental gingivitis (EG). During EG, rapid and consistent alterations in plaque microbiota, metabolites and salivary cytokines emerged as early as 24-72 hours after pause of oral hygiene, defining an asymptomatic ‘sub-optimal health’ (SoH) stage. SoH also features a steep and synergetic decrease of plaque-derived betaine and *Rothia* spp., suggesting an anti-gum-inflammation mechanism by health-promoting microbial residents. Global, cross-cohort meta-analysis revealed a high Microbiome-based Periodontitis Index at SoH state, due to its convergent taxonomical and functional profiles towards those of periodontitis. In contrast, caries SoH features a microbial signature very distinct from caries. Thus SoH is a universal state of polymicrobial inflammations with disease-specific features, which is key to maintaining a disease-preventive plaque.

## Introduction

Gingivitis, the inflammatory lesion of the tooth-supporting soft tissues, is one of the most common oral diseases in humans and has been a global health burden for centuries (*1-5*). It results from a dysregulated immuno-inflammatory response which is induced by dysbiotic plaque biofilm (*6*). Manifested with various clinical signs and symptoms, the gingival condition of gingivitis is affected by both local and systemic factors (*4*). Notably, this inflammatory lesion can be resolved (i.e., reversible) following appropriate professional care. Whereas, uncontrolled gingivitis can progress to the irreversible periodontitis, which is characterized by destruction of tooth-supporting tissues and alveolar bone in susceptible individuals, eventually leading to tooth loss (*7*) and an increased risk of systemic diseases like diabetes and cardiovascular disease (*8-10*). Thus, prognosis and early diagnosis of gingivitis are of great importance in promoting oral health and general well-being (*11*).

However, how gingivitis is initiated remains elusive (*11*). In natural human populations, gingivitis symptoms can be reversible and volatile, as numerous internally or externally imposed disturbances including oral hygiene practices (personal or professional), or impairment of immune system, injury, diet and oral state can all affect disease development and confound disease prevention and monitoring (*12*). Population-wide microbiome associations have unveiled the compositional shifts of plaque during gingivitis progression (*13-17*), and the functional potential of oral microbiome in gingivitis onset was profiled via metagenomics and metatranscriptomic approaches (*15, 18, 19*). However, due to the lack of longitudinal perspective that includes each of the players of microbiota, their metabolites and host immune response, the molecular mechanisms underlying gingivitis onset and progression remain ill defined (*13, 19*).

As for periodontitis, the irreversible and detrimental stage of gum inflammation resulted from chronical, uncontrolled gingivitis, a distinct phylogenetic structure of oral microbiota in diseased hosts versus healthy ones was revealed via 16S rRNA gene or metagenome sequencing (*20-23*). In particular, multiple separate cohort studies have probed the functional potential of periodontitis-associated microbiota via metagenome (*15, 18, 19, 24, 25*) or metatranscriptome (*26, 27*). However, the inherent mechanistic link of gingivitis and periodontitis, which is crucial to clinical prevention and treatment of both diseases, has remained elusive, due to (*i*) the high degree of heterogeneity among host hosts and variation in experimental procedures among the microbiome profiling endeavors, and (*ii*) the inability to track both microbiome and host factor and interrogate their interaction over the full course of gingivitis-to-periodontitis progression within an individual.

To address these key challenges, herein we leveraged a longitudinal, multi-omics experimental design that includes personalized microbial, metabolite and host immuno-response profiles, to provide a high-temporal-resolution, system-level, mechanism-based landscape of the transition from periodontal health, to onset of gum inflammation and eventually to gingivitis (**Fig. 1**). These efforts unveil a microbiome-defined ‘sub-optimal health’ (SoH) stage of gingivitis, at just 24-72 hours after pause of oral-hygiene-practice, which is symptom-free yet carries a microbial signature highly similar to periodontitis. Despite its lack of symptoms, SoH features a steep decrease of microbe-produced betaine, whose abundance is synergetic with *Rothia* spp. yet negatively correlated with bleeding, suggesting an anti-gum-inflammation mechanism by health-promoting residents of plaque. Discovery of this microbiome-defined, symptom-free SoH stage is valuable to prevention and intervention of periodontal diseases. Moreover, meta-analysis of past gingivitis, periodontitis and caries microbiome studies revealed a high Microbiome-based Periodontitis Index for the gum SoH state, yet in contrast, caries SoH features a microbial signature very distinct from caries, suggesting SoH as a shared state of chronic polymicrobial inflammations that carries disease-specific features.

**Fig. 1.**
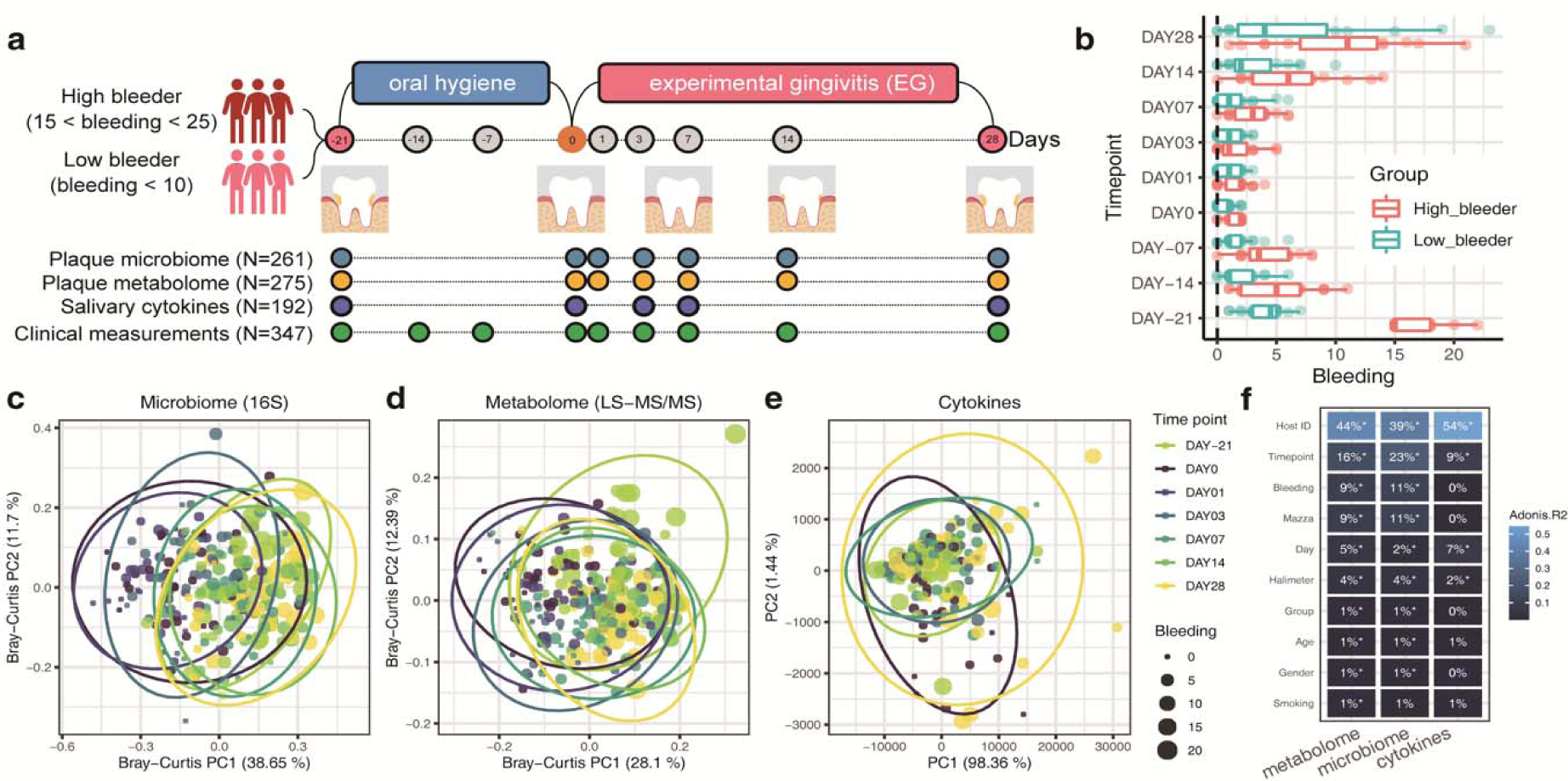
The longitudinal multi-omics landscape of gingivitis onset and progression in a human population. (**a**) Experimental design. Among the 40 healthy adult volunteers that participated, 20 were healthy subjects (with < 10 Mazza bleeding sites), and the rest of them were unhealthy ones (Mazza bleeding sites from 15 to 25) at the start (Day -21 or NG). This study yielded clinical measures (at nine time points), oral microbiome and metabolome data from supragingival plaque samples (at seven time points), and host immune response data from salivary samples (at five time points) for each of the 40 subjects. (**b**) Temporal changes in the clinical symptoms for volunteers. Boxes represent the interquartile range (IQR) and the lines inside represent the median. Whiskers denote the lowest and highest values within 1.5x IQR. (**c and d**) Principal coordinates analysis (PCoA) based on the genus-level Bray–Curtis dissimilarity of (**c**) plaque microbiomes (16S-amplicon sequencing), and (**d**) metabolome profiles (LC-MS/MS); were shown. (**e**) Principal component analysis (PCA) of the salivary cytokine profiles. Each dot in PCoA or PCA represents a plaque or saliva sample and is included in an ellipse whose color indicates time point. Each dot is also sized based on the severity of symptom (gum bleeding). (**f**) Comparing the quantitative variation in all measurements explained by the major factors. PERMANOVA shows that inter-individual variation is the largest factor for all measurement types, while time and disease phenotype also capture sizable variations. Asterisks: FDR-corrected statistical significance (FDR * *p*≤ 0.05, ** *p* ≤ 0.01, *** *p* ≤ 0.001)

## Results

### An experimentally tractable model of gingivitis onset and progression

To control for the many confounding factors (e.g., individuality in initial gum health state or in oral-hygiene behavior) for host–microbiome dysbiosis during gingivitis (i.e., the earlier stage of periodontal disease), we designed for a 40-adult cohort an experimentally tractable model of gingivitis onset and progression (*13, 16*) (**Fig. 1, Table S1**). Specifically, on Day -21 (natural gingivitis, or NG), all 40 adults were randomized into two groups: either high (15 to 25; 20 subjects) or low (0 to 10; 20 subjects) bleeder (**Methods**; **Fig. 1a**). These hosts then underwent a rigorous oral hygiene regimen (dental scaling) for three weeks, resulting in greatly reduced bleeding (median gingival bleeding of 1) on the baseline state of Day 0 (“Baseline”, i.e., a healthy gingival state). Next, the subjects underwent a four-week program inducing experimental gingivitis (EG), which greatly and consistently elevated gingival bleeding, until Day 28 (*p*<0.01 for gingival bleeding; i.e., the diseased state; **Fig. 1b**). Notably, the between-group symptomatic difference at NG (*p*=1e-22, *t*-test), the basis for the high-/low-bleeding stratification of hosts at NG (i.e., Day -21), is much greater than any of the subsequent time points (both before and after Baseline) (**Fig. 1b**). In fact, mild or marginal difference in bleeding between high and low bleeders was observed at the seven subsequent time points (*p*<0.05, *t*-test), but no such symptomatic difference is found at Day 1 or 3 (*p*>0.05, *t*-test). This suggested that disease severity in natural population (i.e., NG) is not necessarily deterministic among individual hosts, and the high-bleeders can recover almost as rapidly and thoroughly as low-bleeders if they follow a proper oral hygiene practice.

Integrated longitudinal profiles of both microbial and host immune programs were unveiled via 275 supragingival plaque (simultaneously for taxonomy and metabolome, via 16S rRNA amplicon sequencing and liquid chromatography-tandem mass spectrometry (LC-MS/MS)-based mass spectrometry, respectively) and 192 matching saliva samples (cytokine profile via multiplexed bead immunoassay) collected by professional dentists. The plaque microbiome, plaque metabolome and salivary cytokines were profiled at time points that fully span the entire 49-day NG-Baseline-EG course (from Day -21 to Day 28 day) while densely sampled the transition from Baseline (0 day) to the onset of EG (e.g., Day 1, 3 and 7; **Fig. 1a, Fig. S1**), so that the tertiary interplay can be temporally monitored, especially at disease onset.

Symptomatic severity (i.e., bleeding) contributed greatly to the first principal coordinate in PCoA of plaque microbiome (**Fig. 1c**) or metabolome (**Fig. 1d**). For plaque-related measurements, although inter-individual variation accounts for majority of symptomatic variance (40-45%; **Fig. 1f**), disease status (9-11%) or time point (16-23%) also explains much of it (**Fig. 1c-d, f**). In contrast, no significant correlation was found between bleeding and salivary cytokine profile (**Figs. 1e-f**). Notably, time point still explains 9% variation in cytokine profile (although the inter-individual factor accounts for 54%) in fact, many salivary cytokines respond to gingivitis development only at the initial time points post Baseline such as Day 3 or 7 but did not further increase afterwards when hosts accumulated even more bleedings. This suggests that the oral host-microbe interplay is the most intensive at the onset stages of gingivitis.

Therefore, we hypothesized the Day 1-3 after dental scaling as the “SoH” stage (**Fig. 1b**). At this stage, we did not detect within-host temporal difference in clinical symptoms (i.e., from Day 1 to 3 after dental scaling) (*p*>0.05, paired *t*-test, **Fig. 2a**), however, the microbiome in the supragingival plaque and even host immune molecules might have dramatically changed due to the detrimental environmental disruptions in EG induction (i.e., poor oral hygiene).

**Fig. 2.**
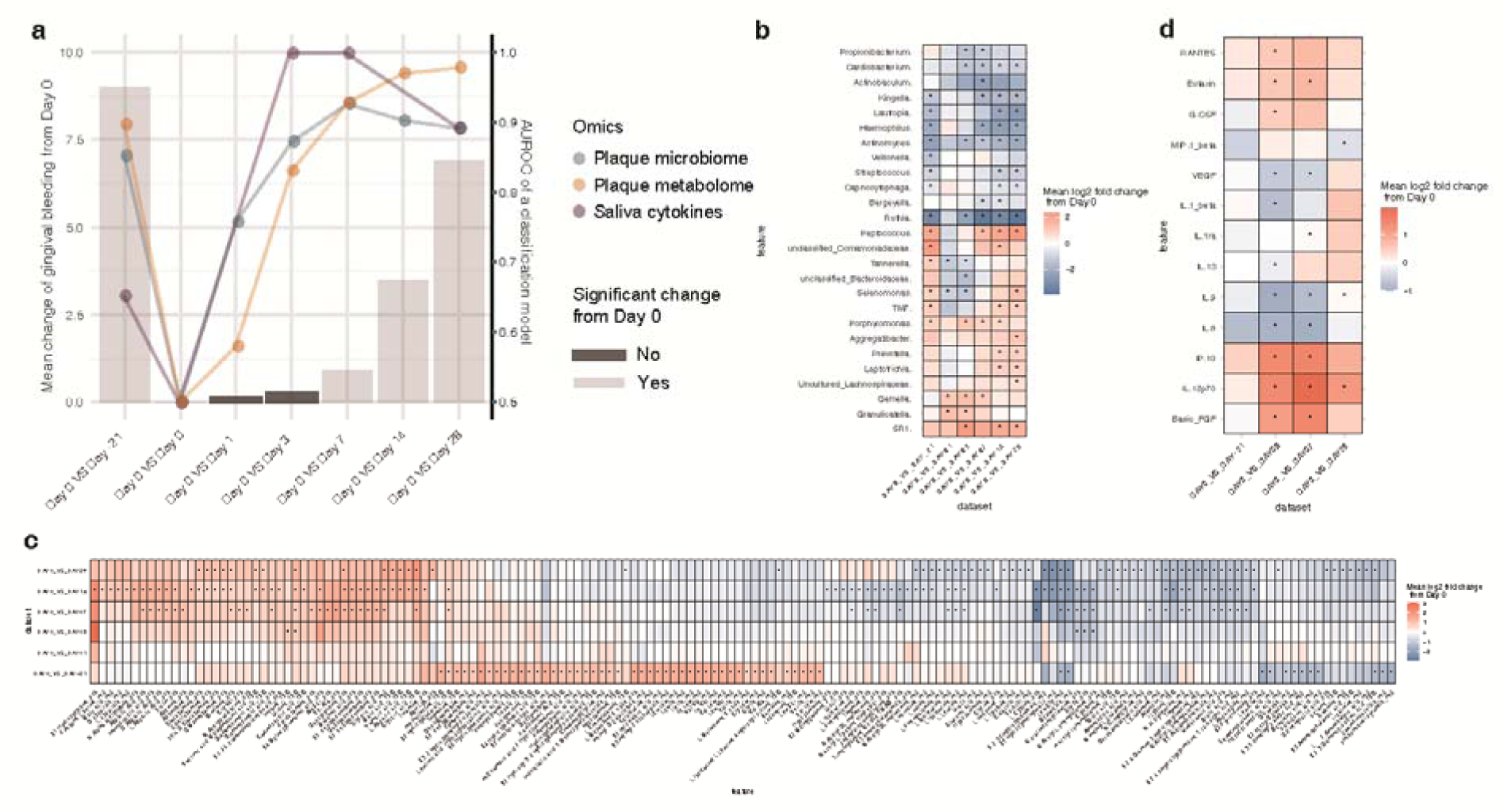
A plaque-microbiome-defined SoH stage that takes place earlier than the emergence of clinical symptoms. (**a**) The symptomatic change (i.e., mean bleeding difference) within hosts (n=40), between each of the time points (Day -21, Day 1, Day 3, Day 7, Day 14, Day 28) and Baseline (Day 0). Color of bars shows FDR-corrected statistical significance: in particular, Day 1-3 are the “SoH” stage when no change in clinical symptoms as compared to Baseline was observed within the hosts. The scatter plots show the AUROC (the *y* axe on the right) of classification models using plaque microbiota, plaque metabolome or salivary cytokines between Day 0 and each of the other time points (Day -21, Day 1, Day 3, Day 7, Day 14 and Day 28). In (b, c and d) we identified molecular features from each measurement type that were differentially abundant at a time point as compared to Day 0. (b) The heatmap for the mean log2 fold changes of microbial responders (with significance threshold Bonferroni *p<*0.05) in plaque during the onset and progression of NG. (**c**) The heatmap for the mean log2 fold change of both early and persistent metabolite responders (with significance threshold Bonferroni *p<*0.05) in plaque. On the *x* axis, “pos”/“neg” after a chemical compound name indicates acquisition via a positive/negative ionization mode in the non-targeted metabolomic approach, while “TSQ” indicates acquisition via from the targeted metabolomic approach. (**d**) Heatmap for the mean log2 fold change of cytokines at each time point (Day -21, 3, 7 and 28) versus baseline (Day 0). Blue denotes reduction while red shows enrichment (versus Baseline). Asterisk: Bonferroni-corrected statistical significance (* *p* ≤ 0.05). No asterisk: no significant change.

### Profound disruption of plaque microbiota/metabolome and salivary cytokines at SoH

To quantitatively measure the shifts in the plaque microbiome and host immunity in the emergence of clinical symptoms, we established a unified metric to measure the temporal changes in multi-omics data from Baseline to EG. Between-timepoints classifiers of host gingival status were built from plaque microbiota, metabolome and salivary cytokine profiles, via the random forests (RF) algorithm. On top of those RF models, we employed a model-accuracy metric (AUROC) as a proxy to quantify the temporal changes of each measurement type at each of the timepoints (i.e., Day -21, 1, 3, 7, 14, and 28) from Day 0. Furthermore, to dissect the multi-omics associations, we compared temporal changes in AUROC values of RF classifiers related to plaque microbiome, plaque metabolome, and salivary cytokines together with those from the clinical symptoms (**Fig. 2a)**. Unexpectedly, the AUROC of RF classifiers for plaque microbiota rapidly shifted in the first 3 days (0.75 at Day 1 and 0.87 at Day 3) from Baseline: it already resembled Day-28 microbiota (severe gingivitis stage; AUROC=0.89) as early as Day 3 (**Fig. 2a**), and actually saturated after Day 3. Therefore, a microbial SoH stage occurred earlier than the emergence of clinical symptoms. In concordance with plaque microbiota, the AUROC on the plaque metabolome increased quickly from 0.58 (Day 1) to 0.92 (Day 7) within 7 days yet did not plateau until after 14 days (AUROC=0.97), suggesting the plaque metabolome was persistently shifting toward a gingivitis-like state. However, the most abrupt changes in the plaque metabolome also took place in the first three days after dental scaling (**Fig. 2a**), indicating that plaque metabolome change also precedes the development of bleeding symptoms, well before they are detectable by professionals. Notably, despite the concordant changes over time between plaque microbiota and metabolome, the saturation of the AUROC of metabolome-based RF classifiers was 7 days later than that of microbiota-based classifiers (**Fig. 2a**), suggesting microbiome-shift dependent changes in the plaque metabolisms during gingivitis onset.

Interestingly, in the SoH stage, the immune response was even more pronounced than both plaque microbiota and metabolome (**Fig. 2a**). The AUROC reached up to almost 0.99 at either Day 3 to 7, while the median gingival bleeding within this period (1 for Day 3 and 2 for Day 7) was relatively low. In contrast, the AUROC at Day -21 (i.e., naturally occurring gingivitis) and Day 28 were all even lower than that in the SoH stage, while the median gingival bleeding was relatively high (8 for Day 28, 11 for Day -21). This suggests that the alterations in the cytokine profiles are not necessarily associated with disease severity but are a response to the intensity or magnitude of organismal and metabolite changes in the plaque microbiome.

The longitudinal concurrent metabolomics and 16S amplicon microbial community profiling from dental plaque samples elucidated the reassembling process of supragingival plaque biofilms after dental scaling (**Fig. 2a**). A key question then is to identify potential microbial and metabolic factors that drive the microbial dysbiosis in the plaque. Thus, to compare the microbiome responses across different stages of disease progression, we performed differential abundance analysis on the CLR-transformed relative abundances of each genus-level taxon between a given time point (Day -21, 1, 3, 7, 14 and 28) and Baseline (Day 0), and compared the results across the stages of EG (Wilcoxon rank-sum test with the Bonferroni correction) (**Fig. 2b**). The microbial markers persistently enriched/depleted with gingivitis progression (such as *Porphyromonas* and *Rothia*), were termed ‘persistent responders’, while those genera transiently enriched/depleted at the early stage of gingivitis progression (i.e., Day 1-3) were ‘early responders’ (such as *Gemella*). Similarly, for plaque metabolome, we identified a series of persistent and early responders in gingivitis development: over 50 metabolites were persistently over-or under-represented during disease development and therefore provided a clue to path-physiology of gingivitis (**Figs. 2a, d**).

Accordingly, time-resolved, differentially abundant cytokines in saliva at Day -21, 3, 7 and 28 were also identified (as compared to Day 0; **Fig. 2c**). Eleven out of the 27 salivary cytokines, such as eotaxin, IL-5, MiP1-beta, IFN gamma, basic FGF, and GSF, altered early, within 72 hours from Baseline (i.e., at the SoH stage), yet did not exhibit any significant difference from Baseline in later, gingivitis-developed timepoints (e.g., Day 28, the most severe gingivitis states along the course). In fact, the SoH stage is featured by a prominent activation of both pro- and anti-inflammatory cytokines that stabilized in later stages of EG (**Fig. 2a, c**). Notably, cytokine alterations are more correlated with particular phases such as SoH than with gingivitis severity, which underscores the importance of high-resolution temporal view of the host-microbiome interplay.

### Integrated microbiome-metabolome dynamic profiles of oral biofilms underlying SoH

To identify plaque microbial activities that underlie gingivitis onset and progression, we constructed a cross-measurement type association network that incorporated both microbial taxa and metabolome from the 261 plaque samples. To reveal trends in the data, Procrustes analysis was used to direct compare the different omics data sets (of identical internal structure) on a single principal coordinates (PC) analysis (**Fig. S3**). Overall, strong correlation between microbial taxa and metabolome of all plaque samples was observed along the NG-Baseline-EG course (*r*=0.53). In fact, such agreement between microbial taxa and metabolomics did not vary with the gingivitis progression (**Fig. S3**), suggesting the key roles of microbes-derived metabolites in this process.

We then built a co-occurrence network from the multi-omics data for biomarker discovery, by calculating the correlation matrix of all features via Spearman’s correlation analysis. The resulting network contained 27,942 total significant edges (|rho|>0.6, FDR *p*<0.05) and 1196 nodes that span features from all three types of measurement. A filtered subnetwork was further built from 29 bacterial genera, 304 metabolites, and 8 salivary cytokines that were differentially abundant between Day 0 and 28 (**Fig. 3a**). Between-metabolites associations accounted for the vast majority (over 99%) of edges, clearly revealing complex and strong association among metabolites. In addition, 51 strong co-associations between microbial genera and metabolites were found, highlighting the impact of gingivitis onset and progression on microbe-dependent metabolisms in plaque. Among these, the *Rothia*-betaine link is one of the most prominent features in the network (red arrows in **Fig. 3a**). As a gingivitis-depleted bacterial marker, *Rothia* had the most links to metabolites (n=14) and exhibited the strongest association with the metabolite of betaine (i.e., trimethylglycine or TMG; rho=0.7; **Fig. 3a**), which is also gingivitis-depleted. In fact, the abundance of betaine and *Rothia* are highly synergic along the full 49-day course (**Fig. 3b**); moreover, both were negatively correlated with symptomatic severity of gingivitis: depleted from NG to Baseline and then enriched again from Baseline to EG, with the peaking of betaine and *Rothia* coincident with the maximal healthy state of gingivae at Baseline (**Fig. 3b**). Notably, the depletion rates of betaine and *Rothia* post during EG induction are not constant: they both steeply decreased during the SoH stage and then gradually stabilized (**Fig. 3b**); in particular, for *Rothia*, at Day 3 its level already dropped to 21% of its peak at Day 0, then it bottomed at Day 7 and stayed so for the remaining 21 days). These observations suggest that the SoH stage, despite the lack of clinically observable changes in bleeding (vs. Baseline), is the most active and consequential phase in both microbiome structural change and the gingivitis-driving microbial metabolism.

**Fig. 3.**
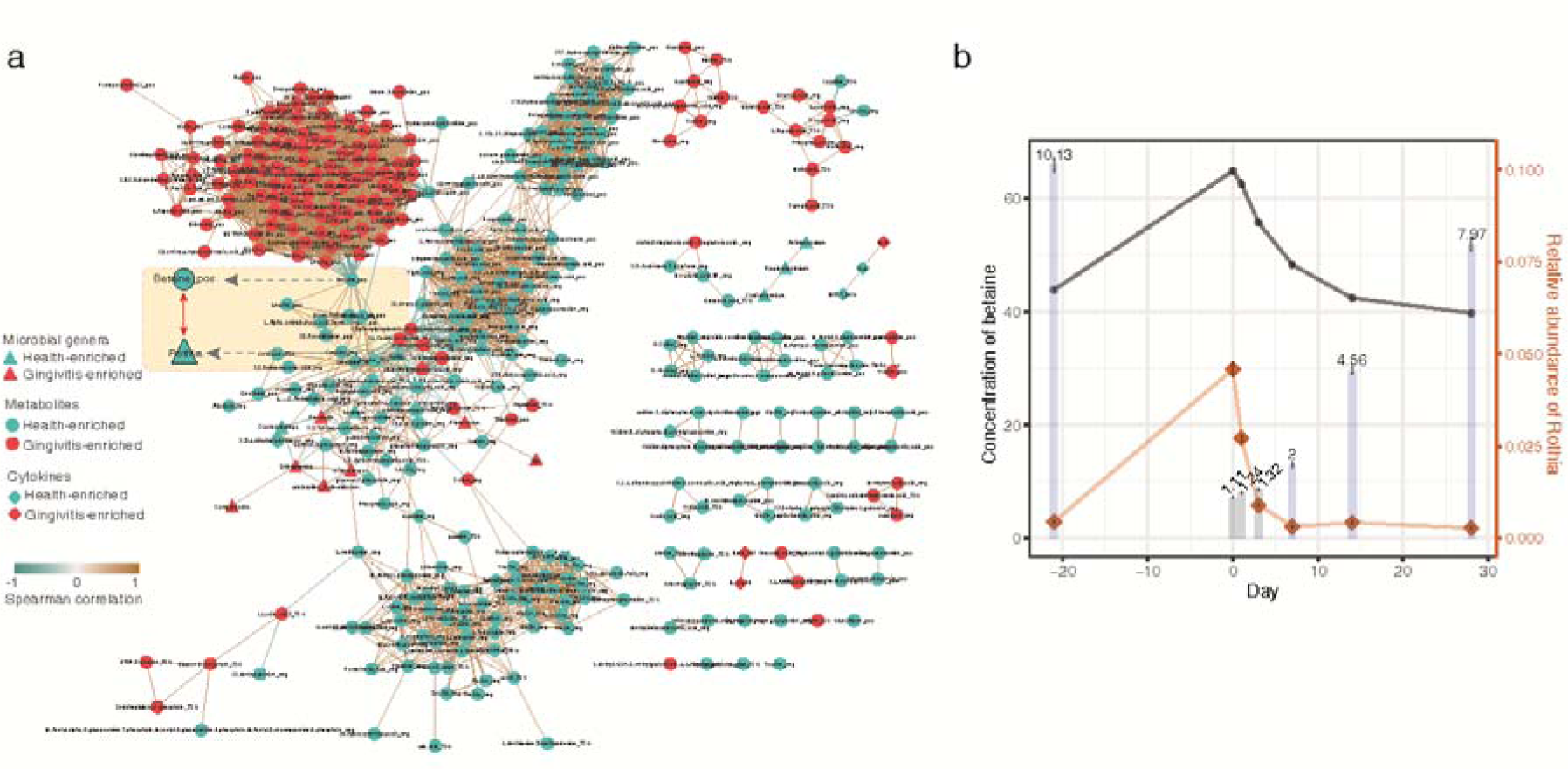
The interplay of plaque taxa, plaque metabolites and salivary cytokines during gingivitis retrogression, onset and progression. (**a**) Network analysis of microbial taxa and metabolites in the temporal program of NG-Baseline-EG. Negative correlations are shown in green, positive in blue and predictive taxa in gray. Edge weights represent the strength of correlation. *Rothia* and betaine have the largest number of connections (i.e., they are the hub nodes) and are highly correlated to each other. For node of metabolites, “pos”/“neg” indicates acquisition by a positive/negative mode in the non-targeted metabolomic approach, while “TSQ” indicates acquisition from the targeted metabolomic approach. (**b**) The temporal co-variation of betaine and *Rothia*, along the process of gingivitis retrogression and induction. The bar plot indicates the clinical symptoms (i.e., mean bleeding) at each of the time points (Day -21, Day 0, Day 1, Day 3, Day 7, Day 14, Day 28). Color of bars shows statistical significance in bleeding between a given time point and Baseline (Day 0): significant (blue) and not significant (grey).

Coincidentally, in addition to its synergy with heath-enriched bacteria such as *Rothia*, betaine is negatively linked to many gingivitis-enriched ones such as *Peptostreptococcus, Prevotella*, and *Treponema* etc (**Fig. 3a**). This suggests an important, perhaps protective, role of betaine in gingival inflammation. Accumulating evidence has shown that betaine plays an anti-inflammatory role in multiple inflammatory diseases, potentially by balancing hyperosmosis and protecting cells from shrinkage and death (*28*). Similarly, the positive link to betaine and the negative association with gingivitis severity indicate that *Rothia* is perhaps beneficial to gingival health and it potentially contributes to betaine metabolism in plaque.

On the other hand, only three out of the 27 cytokines tested are present in the network (**Fig. 3a**). MiP1-beta is enriched in healthy gingivae, yet IL-9 is enriched in gingivitis and negatively correlated with MiP1-beta (**Fig. 3a**): in fact, IL-9 is significantly downregulated at Day 3 and Day 7 and upregulated at Day 28 (versus Day 0; **Fig. 2d**). However, no specific associations between salivary cytokines and plaque taxa or plaque metabolites were found over the process of EG induction (**Fig. 3a**).

### Identifying microbiome links between gingivitis-SoH and periodontitis via meta-analysis

To derive a microbiome-based view of the gingivitis to periodontitis transition (a process that can take decades), we conducted a meta-analysis of published microbiomes for gingival plaques, of sufficient sample size (>20 human adults) and with disease-associated (i.e., case or control labels) or time-revolved metadata (i.e., baseline or time point labels) (**Table 1**). Among the datasets found (all 16S rRNA amplicon based), six were publicly accessible, thus collectively 1505 oral microbiome samples reanalyzed from raw sequences (via Parallel-Meta 3.0 (*29*) and Oral Core microbiota database; **Table 1**; **Fig. 4a, b**), for taxonomic profiles and metabolic functions (via PICRUSt (*29, 30*); **Fig. S5b**).

**Table 1.** The gingival-inflammation microbiome datasets used in the meta-analysis.

| Dataset | Disease related | Sampling niche | Sampling method | Ref. | Target region | Primer | Seq platform | DNA extraction kit | Sample size | Host population size | Host geolocation | Data source |
| --- | --- | --- | --- | --- | --- | --- | --- | --- | --- | --- | --- | --- |
| CN_SoH | Gingivitis | Supragingival plaque | Plaque were collected with sterile Gracey curettes and then removed from the curettes with a cotton-tipped swab. | this study | V1-3 | 5F-534R | Miseq | QIAamp DNA Mini Kit | 261 | 40 | Beijing, China | http://mse.ac.cn/SoH.html) |
| CN_EG_2014 | Gingivitis | Supragingival Plaque | Supragingival plaque samples (along the gingival-line within 2 mm depth) from two entire quadrants (1&3 or 2&4) were collected by sterile Gracey curette at each visit. | (13) | V1-3 | 5F-534R | 454 | Bead-Beating and Lytic-Enzyme-Cocktail Master-Mix were used for bacterial lysis; DNeasy® Blood & Tissue Mini Kits also used. | 150 | 50 | Beijing, China | SRP022235 , SRP022233 |
| CN_EG_CPC_2016 | Gingivitis | Supragingival Plaque | Same as above | (17) | V1-3 | 5F-534R | 454 | Same as above. | 123 | 41 | Beijing, China | SRP022233 |
| CN_AntiG | Gingivitis | Supragingival Plaque | Same as above | (16) | V1-3 | 5F-534R | 454 | Same as above. | 398 | 99 | Beijing, China | SRP045295 |
| UK_EG+Perio<br>dontitis | Gingivitis,<br>Peridontitis | Supragingival and subgingival plaque | Supragingival plaques were collected using a sterile curette from all the mandibular teeth with the exception of the third molars. Subgingival plaques were collected by inserting a curette to the full depth of pockets >6 mm, after the removal of supragingival plaque. | (15) | V1-3 | 27F-519R | 454 | GenElute Bacterial DNA Extraction Kit (Sigma-Aldrich). | 92 | 20+20 | London, UK | SRP026653 |
| US_Periodontitis | Periodontitis | Subgingival plaque | After removing supragingival plaque and drying the target sites, subgingival samples were collected by insertion of four medium paper points for 10s into three sites. Deep and shallow sites were sampled separately from subjects with periodontitis. | (23) | V1-2; V4 | 27F-342R | 454 | QIAamp DNA mini kits | 87=29(periodontitis shallow)+29(periodontitis deep pocket)+29(Health) | 58 | USA | SRP009299 |
| ECC | Early childhood caries | Subgingival plaque and saliva | Dental plaques collected from all erupted deciduous teeth by brushing for 1 min via a sterile toothbrush. Unstimulated saliva produced during 5 minute was collected in 50 ml sterile tubes. | (31) | V1-V3 | 5F-534R | 454 | The MO BIO PowerSoil DNA Isolation kit with minor modifications | 284 | 50 | Guangzhou, China | SRP04094, SRP040947 |

**Fig. 4.**
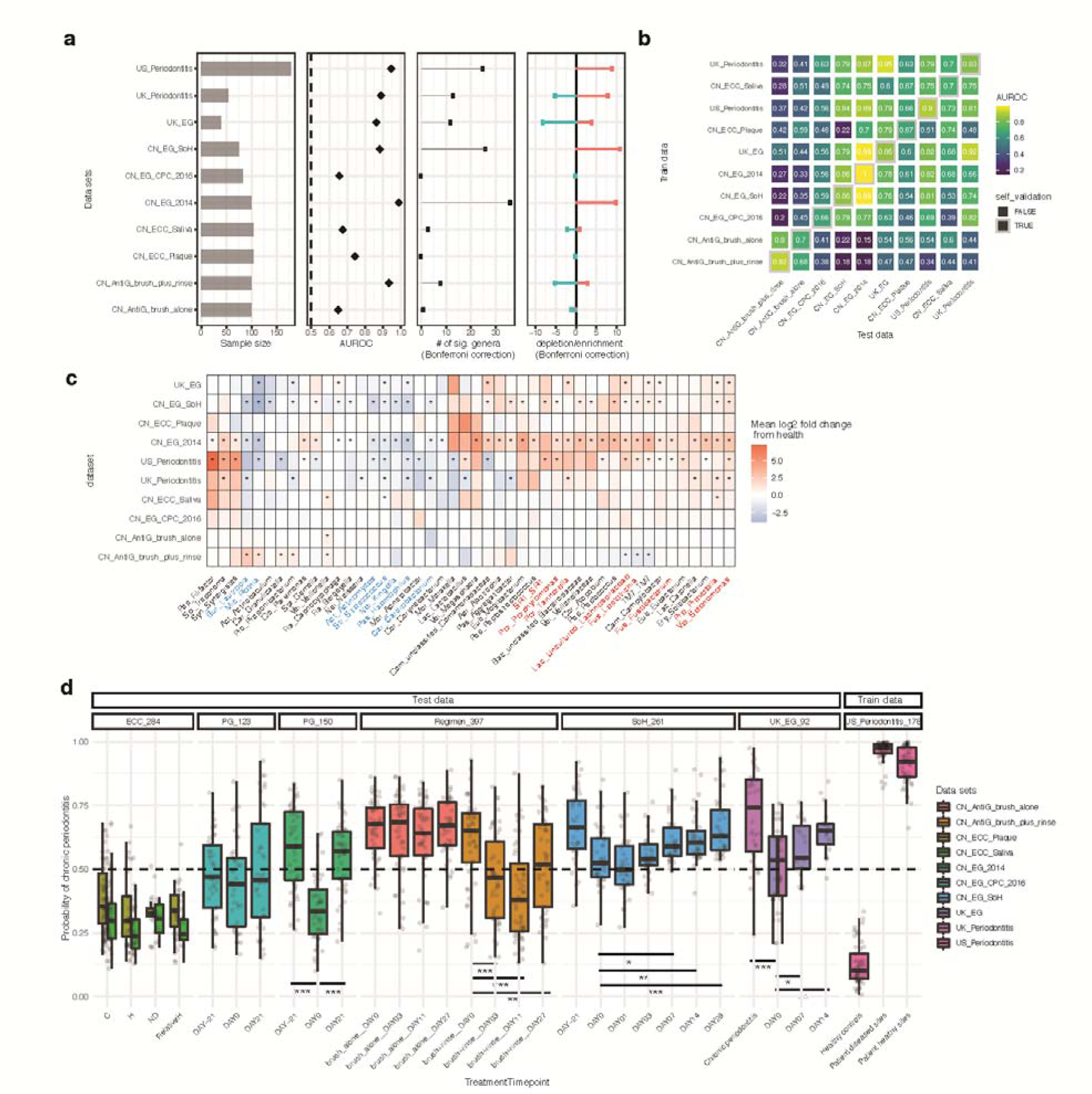
Meta-analysis of existing gingival microbiome datasets revealed similar microbial signature between gingivitis-SoH and periodontitis. (**a**) Most periodontal disease progression or retrogression show microbiome alterations, with consistent disease-associated shifts that differ in their extent and direction. Panels from left to right: (*i*) sample size for each study; (*ii*) area under the ROC curve (AUROC) for the genus-level random forest classifiers (X-axis starts at 0.5, the expected value for a classifier that assigns labels randomly, and AUROCs < 0.5 are not shown); (*iii*) number of genera with *q* < 0.05 (Wilcoxon rank-sum test, Bonferroni correction) for each data set (if a study reveals no significant associations, no points are shown). (*iv*) direction of the shifts in microbiome structure, i.e., the percentage of associated genera that are enriched in disease. (**b**) Cross-prediction matrix reporting prediction performance as AUROC values obtained using a random forest model on the genus-level relative abundance. Matrix values refer to the AUROC values obtained by training the classifier on the dataset of corresponding row and then applying it to the dataset of corresponding column. The prediction accuracy between gingivitis and periodontitis is remarkably high, suggesting a strong microbial link between these two periodontal diseases. Moreover, the prediction accuracy between anti-gingivitis treatments is higher than that between EG experiments, suggesting anti-gingivitis treatments often result in very similar microbiome responses, regardless of the difference in cohorts. (**c**) Heat map for log2 mean fold change of all plaque genera between the last day of treatments and Baseline in each of the longitudinal studies (or between case and control groups in the cross-sectional studies). Blue denotes reduction in relative abundances of genera (red: enrichment) versus Baseline. Those significant fold changes (Bonferroni-corrected *p*<0.05) are marked by asterisks, while not-significant fold changes (Bonferroni-corrected *p*>0.05) are indicated as blank in the heatmap. Text color of the genus names indicates those showing highly consistent enrichment (red) or reduction (blue) in the periodontal disease state across data sets. (**d**) A Random Forests classifier of periodontitis was built based on the subgingival microbiomes in a US periodontitis cohort, and then applied to all the other datasets in the meta-analyses, so as to model the estimated probability of periodontitis for the gingivitis patients. Asterisks: FDR-corrected statistical significance (FDR * *p*≤ 0.05).

We first tested whether the reported microbiome associations with the oral disease states or the anti-gingivitis treatments can be recapitulated (**Table 1**). To compare across studies such disease-responses of microbiome, we first grouped all data into ten “datasets”. Each dataset can include samples from case and control groups in a cross-sectional study (e.g., “UK_Periodontitis”) or samples at the baseline and subsequent time points in a longitudinal study of EG (such as “CN_EG_2014”) or an anti-gingivitis treatment (such as “CN_AntiG_brush_plus_rinse”). Next, for each dataset, we built a genus-level RF classifier to distinguish disease states (gingivitis, periodontitis, or dental caries) from the health states longitudinally or cross-sectionally, and then compared their AUROC across datasets.

Surprisingly, periodontal disease status can be classified between hosts or within hosts (AUROC>0.7) in all studies (**Fig. 4a**). Notably, the states of gingivitis or chronic periodontitis are highly distinguishable by plaque microbiome (AUROC>0.9) in six out of eight related datasets (**Fig. 4a**). We then asked whether and to what extent the microbiome-based RF classifiers of periodontal disease states can be applicable from one dataset to another (**Fig. 4b**). For gingivitis, we observed very limited degradation in prediction accuracy for the cross-trained RF models from one cohort to another (AUROC ranges from 0.88 to 0.99 in either self-validation or prediction). Moreover, a RF classifier trained on periodontitis can be readily applicable to gingivitis or vice versa (AUROC>0.75 in either self-validation or prediction), despite the large technical difference (or other non-disease-related biological differences) between studies/cohorts in the microbiome data that frequently confound such cross-applications (**Fig. S4a**). Thus, the gingivitis and periodontitis classifiers share a large number of microbial markers, suggesting a high degree of similarity in the underlying microbiome.

Then, the microbial signatures associated with gingivitis or periodontitis were compared across these datasets (**Methods**). *Firstly*, we asked whether the identified microbial response to gingivitis onset (i.e., SoH) or progression is consistent with reported gingivitis microbiome in these independent cohorts. Here 1023 samples (N=931 from China; N=92 from UK) from five gingivitis-related datasets were compared, each with a longitudinal design that tracks microbiome dynamics along gingivitis progression or retrogression. For cross-study comparison of microbial responses, statistical analyses on samples from the baseline and the last time point in each study were performed (with univariate tests on genus-level CLR-transformed relative abundances conducted for each dataset independently and the results compared across studies; Wilcoxon rank-sum test with the Bonferroni correction). Notably, the gingivitis-associated microbiomes are highly reproducible across studies (**Fig. 4c**). In the EG datasets, microbiome shifts are characterized by enrichment of a large proportion of ‘pathogenic’ or pathogen-associated genera and depletion of a few commensal oral bacteria (consistent across studies; **Fig. 4c**). The EG-associated microbiome identified from our previous study (i.e., “CN_EG_2014”) harbored the broadest spectrum of microbial shifts (N=41), among which >60% of microbial markers (e.g., *Rothia, Haemophilus, Actinomyces, Streptococcus, Selenomonas, Prevotella, Leptotrichia*, uncultured *Lachnospiraceae*, and TM7) actually overlapped with those identified in other gingivitis-progression studies (including the present gum SoH study; **Fig. 4c**). Moreover, the two anti-gingivitis treatments of brush alone and brush plus rinse(*16*)) are both characterized by enrichment of health-associated bacteria yet depletion of ‘pathogenic bacteria/; in fact, the microbial taxa shifted toward the healthy state during gingivitis retrogression have largely overlapped with markers of the EG studies (e.g., *Lautropia, Rothia, Granulicatella*, TM7 and *Leptotrichia*; **Fig. 4c**), yet in exact opposite directions of abundance change,

*Secondly*, we tested whether or to what extent the stage-specific plaques of gingivitis are linked to those of periodontitis. Specifically, 260 samples were collected from two case-control studies (UK, N=92; US, N=178) on periodontitis microbiome: the UK_Periodontitis dataset where Kistler et al. profiled plaque microbiome of chronic periodontitis (*15*) and the US_Periodontitis dataset where Griffen et al. compared subgingival plaque microbiota from 29 periodontally healthy controls and 29 subjects with chronic periodontitis (including periodontally healthy and diseased sites) from a US cohort (*23*). Notably, the periodontitis microbiomes feature a large number of genera that overlap with those identified in the EG or even the SoH stage of gingivitis (**Fig. 4b-c**; **Fig. S4**). The microbiome shifts responding to chronic periodontitis in the US or UK cohorts were characterized by an enrichment of gingivitis-enriched genera (such as *Porphyromonas, Leptotrichia, Selenomonas*, TM7, *Prevotella*, uncultured *Lachnospiraceae, Campylobacter, Fusobacterium* and *Tannerella*) and a depletion of gingivitis-depleted ones (such as *Rothia, Haemophilus, Actinomyces, Streptococcus* and *Kingella*). Importantly, those gingivitis-associated microbes were all identified as so in the Chinese cohorts. Considering the potential heterogeneity between cohorts (i.e., geographic locations) or technical inter-study batch effects (such as 454 vs. Illumina sequencing platform, different primer sets etc.), the very limited variation in microbial response to periodontal diseases across the two UK/US periodontitis cohorts and the China gingivitis cohort is remarkable.

To validate the similarity in microbiome signature between gingivitis and periodontitis, we built a RF classifier of the chronic periodontitis on the plaque microbiome, and applied this model to a given sample from any of the gingivitis stages for estimating its microbiome-based probability of periodontitis (which we proposed as “Microbiome-based Periodontitis Index” or MPI; **Fig. 4d**). In the training dataset (i.e., US_Peridontitis), MPI of the healthy controls are on average only 10%, while reach up to 99% averagely in periodontitis patients. In our present study, MPI increase progressively along the EG process, a pattern that is consistent with the other EG datasets. In particular, MPI at Day 7 (end of the gum SoH stage), with a median at ∼62%, is significantly higher than that at Day 0, suggesting the emergence of a periodontitis-like microbiome at this stage, due to the aforementioned, profound changes in plaque microbiome, plaque metabolome and host immunity that take place at SoH.

### Comparing microbiome dynamics in the development of gum inflammation and caries

Next, we put the temporal microbial shifts along gingivitis development in a broader context that includes not just periodontitis but dental caries, via meta-analysis of the SoH, UK_EG, UK_Periodontitis, US_Periodontitis and early childhood caries (ECC) datasets(*31*). We classified the disease or pre-clinical status using RF models based on either the species-level taxonomic profile or the predicted functional profile (by PICRUSt) along stages of disease development in all studies (**Fig. 5a**). Surprisingly, AUROC of species-level-taxonomy based RF classifiers for plaque at Day 3 reached 0.85 (function-based classifiers: 0.81), which is already quite close to the 0.88 at Day 28 (function-based classifiers: 0.85). Thus plaque functional profiles already resembles that of the severe gingivitis stage within 24 hours after dental scaling (**Fig. 5a**), and actually saturates after 24 hours. The discriminative power of this function-based classifier (AUROC=0.78) is nearly equivalent to that distinguishing chronic periodontitis patients from healthy individuals from the UK cohort (AUROC=0.82; DAY0_VS_DD), suggesting an ultra-rapid assemblage of functional components in the plaque biofilm that highly resemble those in periodontitis patients. In contrast, in ECC development, oral microbiome did not show as pronounced changes in the early stage (AUROC=0.52; H VS RelativeH) as those in the late stage (AUROC=0.68; H VS C; **Fig. 5a**).

**Fig. 5.**
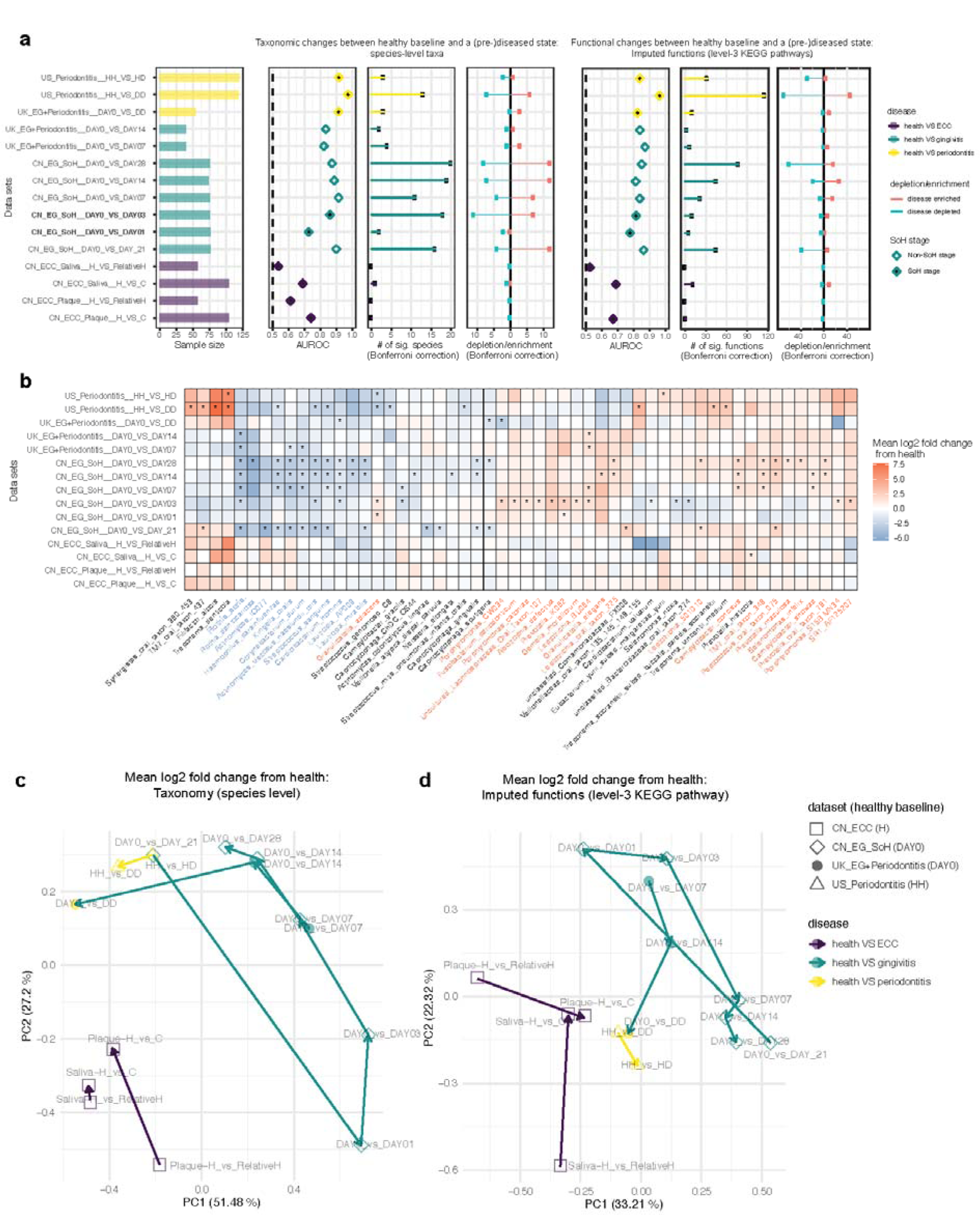
Comparing temporal microbial shifts along disease development between periodontal diseases and caries. (**a**) Most oral disease progression show microbiome alterations, with consistent disease-associated shifts that differ in their extent and direction. Panels from left to right: (*i*) sample size for each study; (*ii*) area under the ROC curve (AUROC) for the species-level RF classifiers (*x*-axis starts at 0.5, the expected value for a classifier that assigns labels randomly; those with AUROCs < 0.5 are not shown); (*iii*) number of species with *q* < 0.05 (Wilcoxon rank-sum test, Bonferroni correction) for each data set. (*iv*) direction of the shifts in microbiome structure, i.e., percentage of associated species that are disease enriched. (*v*-*vii*) Similar analysis conducted on the imputed functional profiles from 16S rRNA sequencing data. (**b**) Heat map for log2 mean fold change of bacterial species between a (pre-)diseased state and the healthy baseline in each. Blue denotes reduction in relative abundances of species (red: enrichment) versus Baseline. Significant fold-changes (Bonferroni-corrected *p*<0.05) are marked by asterisks, while insignificant fold-changes (Bonferroni-corrected *p*>0.05) as blank in the heatmap. We next performed PCoA based on the mean log2 fold change data of species (**c**) or predicted functional pathways (**d**) that are associated with two oral diseases. Each dot in the PCA plots represents a process of microbiome alterations from health to the onset or progression stage of a given oral disease. The dots are colored by diseases. The lines with arrows represent the path that microbial alterations occurred along the disease development.

Moreover, to test whether microbiome successions are concordant between the developmental stages of these oral chronic inflammations, we quantitatively compared the microbial differential abundance profiles between time points or disease severities. For each dataset, the differential abundance (i.e. mean log2 fold change) of microbial features in the plaque/saliva microbiome from healthy baseline to a given developmental stage of disease was measured (**Fig. 5a-b, Fig. S5a**). For two given microbial signatures (e.g. Day0 vs. Day -21 and Day0 vs. Day 28 in the SoH study), we first ranked the features by the degree of differential abundances in each of them and then calculated the Pearson’s correlation between these two feature ranking lists. To reveal the patterns driving the temporal difference in microbiome across diseases, we next performed PCoA via the correlation-based distance metric of all pairs of feature ranking lists, with each dot in PCoA corresponding to a pattern of microbial alteration between the healthy baseline and a particular disease developmental stage (instead of a microbiome sample; **Fig. 5c-d**).

Intriguingly, at the species level, the microbiome differences along gingivitis development are more pronounced than those from periodontitis or dental caries (**Fig. 5c**). During gingivitis progression, along PC1, the profile of microbiome alteration between the baseline (Day 0) and a given time point would increasingly resemble that between health and periodontitis in either the US or the UK cohort. Notably, the microbial taxonomical response to severe gingivitis (e.g. Day0 vs. Day -21, and Day0 vs. Day 28 in the SoH study) is highly similar to that of chronic periodontitis. Thus, taxonomic perturbations during dysbiosis are highly consistent between gingivitis and chronic periodontitis, while the taxonomic responses to the periodontal diseases and dental caries are quite distinct (**Fig. 5c**).

Notably, during gingivitis development, functional potential of microbiome is relatively conservative over time, particularly after the SoH stage (**Fig. 5d**). In fact, our results suggest that the gingivitis-associated community in dental plaque biofilm actually assembles rather rapidly in the very early stage (i.e., the SoH stage), to form a “climax”-like community configuration that is very similar to the periodontitis-associated community (**Fig. 5d**). In contrast, ECC-associated microbiomes at the onset stage (i.e., SoH) are actually very distinct from those at the late stage (*31*). As ECC develops, the primary oral microbial communities (i.e. health-associated) evolves to a convergent state, due to selection of a changed microenvironment of teeth (such as acidification (*32*)), and such a “climatic” state that corresponds to a reliable caries stage, is very distinctive from that in the “new onset” stage of caries (i.e., RelativeH, when no clinically detectable symptoms are apparent in teeth (*31*)) in terms of taxonomic composition or functional profile (**Fig. 5a, c, d**). For example, the cariogenic pathogen of *Streptococcus mutans* are highly enriched in the climax community, yet hardly present at the new onset (i.e., SoH) stage (*31*); in contrast, at the SoH stage of ECC, *Prevotella spp*. exhibit a much stronger statistical power in predicting caries onset than *Streptococcus mutans* (*17*). Therefore, the distinct temporal patterns of microbial succession in plaque-induced pathogenesis, as well as their distinct rates of microbiome change relative to symptom development, appear to be a common stage of such chronic, polymicrobial inflammations that carries disease-specific features.

## Discussion

Despite the technological challenges, integrating the human dental plaque microbiota and metabolomics profiles enables an in-depth and mechanistic understanding towards periodontal disease etiology. Simultaneous analysis of dental plaque samples via DNA sequencing and LC-MS/MS has been hindered by (*i*) the low biomass of dental plaque sampled with high temporal resolution from each host and (*ii*) the difficulty to reconcile the distinct sample preprocessing procedures for DNA sequencing and LC-MS/MS on a plaque sample (e.g., the organic solvent extraction in LC-MS/MS can reduce the DNA quality for sequencing). Therefore, in our new strategy, two dental plaque samples (up to 14 teeth each) were collected (for each subject) from 1 and 3 (plaque A) or 2 and 4 quadrants (plaque B) for sequencing and LC-MS/MS respectively (randomly assigned, to eliminate potential bias). This is particularly enabling for recording the integrated metagenome-metabolome choreography of plaque, when sampled at high temporal resolution, and particularly during the SoH phase (just 0∼3 days away from Baseline, with especially low plaque biomass).

The link and distinction temporal dynamics among host symptoms, immune factors, plaque structure and plaque metabolome unveiled how plaque microbiota drove gingivitis onset and progression. Most importantly, an asymptomatic “SoH” state of gingivae, from 0 to 3 days after dental prophylaxis and pause of oral hygiene, was uncovered, when actually the most intense host-microbiome interactions take place, i.e., rapid and consistent alterations in plaque microbiota, metabolite pool and salivary cytokines. In particular, during this pre-clinical-symptom, very transient gingival state of SoH, plaque residents (e.g., *Rothia* spp.) and metabolites (e.g., betaine) that are strongly negatively correlated with gum-bleeding (over the entire 49-day NG-Baseline-EG process) undergo a steep decrease, while at least eleven salivary cytokines dramatically change in response (six up-regulated and five down-regulated as compared to Day 0) and then rapidly plateau. In contrast, such alterations were not seen in subsequent phases of gingivitis development (e.g., from Day 7 to 28), even for those with much higher symptomatic severity.

Betaine was not previously linked to gingivitis development, despite its being recognized as maintaining cell osmotic pressure which can promote cell survival under the high hyperosmotic pressure potentially due to inflammation and diseases (*28*). Interestingly, it is at present an ingredient in toothpaste for relieving dry mouth (*33*). In our plaque samples, betaine consistently and continuously declined as the gingivitis developed (particularly in the SoH stage), suggesting a protective role against gum inflammation. Notably, its concentration in the plaque was highly correlated with healthy-gum-enrich and gingivitis-depleted plaque residents such as *Rothia* spp.. Therefore, the health-associated members of plaque might have served as a source of betaine that possibly to protect the gum from gingivitis, which underscores the importance of maintaining a healthy plaque.

Notably, although taxonomic shift in plaque took place as early as 24 hours after dental prophylaxis (by acquiring microbial colonizers from saliva (*11, 20*)), it was accompanied by a delayed functional shift as revealed by plaque metabolome. This suggests that establishment of primary colonists in plaque altered within 48 hours (i.e., at or by Day 3) the plaque metabolome, which then elicits both gingival inflammation and subsequent plaque development, starting a detrimental cycle: periodontal tissue destruction by plaque dysbiosis provides nutrients for bacterial growth, which further promotes dysbiosis and tissue inflammation (*11*). Therefore, despite its apparent Baseline-like symptom, the SoH phase is a transient yet crucial time window to prevent or abolish the start of such vicious cycles.

Surprisingly, the implication of this SoH stage finds support from our meta-analysis of past oral microbiome studies, which reveals a microbiome-mediated link between the very early (i.e., SoH of gingivitis) and very late stage (periodontitis) of the periodontal disease which can span decades and affects over half of the global population. Gingivitis and periodontitis patients can share a significant number of bacteria genera (*18-20, 23*), and periodontal treatments can result in depletion of disease-associated bacteria and enrichment of health-associated ones in plaque (*16, 17, 34*). However, systematically tracking microbial associations across different stages for chronic periodontal diseases remains a challenge, since it is impractical to create or modulate advanced disease states directly in humans, while clinical studies can only induce mild or moderate disease states (notably, this holds true for many chronic diseases). Moreover, technical variations such as inter-study differences in the sequencing protocol, 16S databases or statistical methods prevent comparing microbial associations across studies (*35*). For example, microbiome data are compositional (*36*), however in many past studies, traditional statistical methods such as *t*-test or Wilcoxon rank-sum test were widely and inappropriately used on the raw abundance data for microbial marker discovery; in fact, once accounting for the compositionality issues in statistical analysis, it is far less clear whether the reported microbial associations can be recapitulated (*36*).

To tackle these issues, we re-analyzed from raw data all published and accessible microbiome datasets with consistent parameters and RF models. Our results profoundly relate gingivitis to periodontitis via plaque microbiome. Specifically, (*i*) the oral microbiome responses to a disease state, either gingivitis or periodontitis, can be highly consistent across human populations, while this is not the case for most of the other chronic diseases (*31, 35*); (*ii*) the plaque residents specifically responding to periodontal inflammation are quite consistent between the very early stage of gingivitis (i.e., SoH) and the eventually irreversible and detrimental state of periodontitis, despite their decade-long temporal gap and the large host-or technology-related variation among cohorts/studies. This is in contrast to early childhood caries (ECC), where plaque microbiomes at the new onset stage are very distinct from that of the late stage. The patterns and nature of such microbiome change underlying chronic disease development, whether conserved or divergent among the many chronic inflammations in oral or other human body sites, can shed new light on disease etiology and help precise diagnosis, prevention and treatment.

In summary, by tracking the choreography of plaque microbiome structure, plaque metabolome and host immune-response during gingivitis onset and progression, we unraveled a microbiome-defined SoH stage of gingivitis, i.e., the just 24-72 hours after pausing oral hygiene. Although transient and asymptomatic, SoH is a crucial phase when the most intensive changes in plaque structure and metabolism as well as host immune factors take place, and carries a microbial signature highly similar to periodontitis. In light of the epidemic of periodontal disease (*1-5*) and the insufficient public health awareness on oral hygiene (a significant portion of world population still fails to brush teeth daily), our findings underscore the importance of intervening at the SoH stage of gingivitis via proper oral hygiene practices, so as to maintain a healthy, periodontitis-preventative plaque. In addition, since SoH appears to be a shared stage that carries disease-specific microbial, metabolomic and immunological features, it would be promising to define and compare the SoH states of additional chronic polymicrobial inflammations, which should lay the foundation for exploiting their uses in predictive and personalized medicine.

## Materials and Methods

### Overall design of the study

The ‘experimental gingivitis’ notion was established as a non-invasive model in humans for the pathogenesis gingivitis (*13*). This single-center, examiner-blind, controlled clinical trial was conducted at Procter & Gamble (Beijing) Technology Co., Ltd. Oral Care Department, with approval from the P&G Beijing Technical Center (China) Institutional Review Board and in accordance with the World Medical Association Declaration of Helsinki (1996 amendment). ICH Guidelines for Good Clinical Practice (GCPs) were followed. All participants gave written informed consent prior to the study.

### Overview of human cohort

A total of 40 volunteers who met all inclusion criteria participated in this study and all completed it (**Table S2**). Clinical examination of gingival tissues using Mazza index (reference) was conducted at all of the visits by a qualified dental examiner (**Fig. 1a**). For each subject, supragingival plaque and salivary samples were collected by professional dentists at Day -21 (NG), Day 0 (Baseline), Day 1 (EG), Day 3 (EG), Day 7 (EG), Day 14 (EG) and Day 28 (EG), in a longitudinal manner (**Fig. 1a**). The optimal gingival health state on Day 0 was achieved through dental prophylaxis and rigorous oral hygiene during the oral hygiene phase prior to Baseline. Dental prophylaxis including super and subgingival whole-mouth cleaning on a total of 28 teeth was performed on Day -21, Day -14, and Day -7. Subjects were instructed to brush with a sodium fluoride dentifrice three minutes each time twice daily in the oral hygiene phase. On the contrary, in the EG phase from Day 0 to Day 28, only rinsing with purified water was allowed for each of the subjects.

### Clinical assessment

A qualified dental examiner performed oral tissue assessments on the study participants at Day -21, Day -14, Day -7, Day 0, Day 1, Day 3, Day 7, Day 14 and Day 28. Assessment of the oral soft tissue is conducted via a visual examination of the oral cavity and perioral area. The structures examined include the gingiva (free and attached), hard and soft palate, oropharynx/uvula, buccal mucosa, tongue, floor of the mouth, labial mucosa, mucobuccal/mucolabial folds, lips, and the perioral area. Assessment of the oral hard tissues was conducted via a visual examination of the dentition and restorations. Gingivitis was assessed based on the Mazza Index (*13*): sampling was performed on the mesiofacial and the distolingual of each tooth, for a maximum of 56 sites.

### Saliva sample collection

At the Day -21, Day 0, Day 1, Day 3, Day 7, Day 14, Day 28 visits, subjects were asked, prior to plaque sampling, to expectorate approximately 10 mL of unstimulated saliva into a labeled tube (**Fig. 1a**). The samples were frozen at -20°C immediately after collection until use for cytokine profiling.

### Plaque sample collection

Gingival plaque from each of the 40 subjects was collected at Day -21, Day 0, Day 1, Day 3, Day 7, Day 14 and Day 28 (**Fig. 1a**). Specifically, subjects were refrained from oral hygiene practice include tooth brushing, flossing or mouth rinsing in the morning of sampling and supragingival plaque samples along the gingival margin were collected after GI examination using a gracey curette by a qualified dentist. At each time point, to ensure sufficient amount of plaque for analysis, samples were taken from each subject’s maxillary right and mandibular left quadrants or maxillary left and mandibular right quadrants alternatively. All samples were stored under -70°C until use.

### Plaque microbiome structure analyses

Genomic DNA was extracted from the plaques. Barcoded 16S rRNA amplicons (V1-V3 hypervariable region) of all the 261 samples were sequenced via Illumina Miseq. All 16S rRNA raw sequences were pre-processed following the standard QIIME (v.1.9.1) pipeline (*37*). Downstream bioinformatics analysis was performed using Parallel-Meta 3 (*29*), a software package for comprehensive taxonomical and functional comparison of microbial communities. Clustering of OTUs was conducted at the 97% similarity level using the OralCore database (*38*). Taxonomically assigned sequences were further agglomerated at the genus level for structural comparison of microbiomes.

### LC-MS/MS data acquisition for plaque metabolome

Prior to LC-MS/MS analysis, plaque samples were prepared using the following procedures. For extraction, 1 mL 40:40:20 (in volume) MeOH/ACN/Water was added to the pre-weighted supragingival plaque in 2 mL PP tube and vortexed for 1 minute. Plaque pallets in the extraction solvent were incubated in 95°C water bath for 1 hour and then centrifuged at 3000rpm and subsequently transferred to another 2 mL PP tube. For complete extraction, 500 μL extraction solvent was added as described above into the original tube and then vortexed for 10s and centrifuged at 3000rpm for 10 minutes. Each of the final extraction solutions was combined with the other obtained in the last step. Each liquid extraction was dried completely with nitrogen and then stored in -80°C freezer until use.

Non-targeted metabolomic analysis was performed using Q Exactive orbitrap (Thermo, CA). After resuspension of the dried extract, each of the samples (1uL supernatant) was loaded to normal phase chromatography column, then eluted to the orbitrap mass spectrometer with an aqueous phase containing 5mM ammonium acetate as eluent from 1% to 99% within 15 min. The stationary phase was 95% acetonitrile with 5mM ammonium acetate. Data with mass range m/z 100-1500 was acquired at the positive ion mode using data dependent MS/MS acquisition. The full scan and fragment spectra were collected with resolution of 70,000 and 17,500 respectively. The source parameters are as follows: spray voltage: 3000v; capillary temperature: 320°C; heater temperature: 300°C; sheath gas flow rate: 35; auxiliary gas flow rate: 10. Metabolite identification was based on Tracefinder search with home-built database containing 529 compounds.

Targeted metabolomic experiments were performed on TSQ Quantiva (Thermo, CA). C18 based reverse phase chromatography was utilized with 10mM tributylamine, 15mM acetic acid in water (pH ∼6) and 100% methanol as mobile phase A and B respectively. This analysis focused on TCA cycle, glycolysis pathway, pentose phosphate pathway, amino acids and purine metabolism. A 25-minute gradient from 5% to 90% mobile B was used. Positive-negative ion switching mode was performed for data acquisition. Cycle time was set as 1 second and totally 138 ion pairs were included. The resolution for Q1 and Q3 are both 0.7FWHM. The source voltage was 3500v for positive and 2500v for negative ion mode. Sweep gas was turned on at 1(arb) flow rate.

### LC-MS/MS data analysis for plaque metabolome

For targeted metabolomics, triple quadrupole mass spectrometer (TSQ Quantiva, Thermo) was used for the analysis in MRM mode. All the ion transitions and retention times were optimized using chemical standards. Tracefinder (Thermo, USA) was applied for metabolite identification and peak integration. The peaks were manually checked for the analysis. Pooled QC samples were inserted in the batch to ensure system stability.

For untargeted metabolomics, orbitrap mass spectrometer (QExactive, Thermo) was used for the analysis in DDA mode. An in-house database containing MS/MS spectra of over 1500 metabolites was incorporated for metabolite identification. Tracefinder (Thermo, USA) was used for metabolite identification based on MS/MS fragment matching. LS score was applied to confirm the confidence of metabolite identification. Only the metabolites with LS score > 30 were considered as confident confirmation. Otherwise, they were assigned as putative identification. The peaks were manually checked for the analysis. Pooled QC samples were inserted in the batch to ensure system stability.

Normalization was performed before statistical analysis. The missing values were replaced with half of the minimum values in all the samples. Peak areas were normalized relative to the mean of the total area of a sample. Both targeted and untargeted metabolomics data were combined and imported into the R software (version 3.6.2) for multivariate analysis.

### Quantification of salivary cytokines using multiplexed bead immunoassay

We collected 194 salivary samples at Day -21, 0, 3, 7 and 28 from 40 subjects who were selected for quantification of inflammatory cytokines (**Fig. 1a**). All samples were sub-packed (1.0 mL sample in 1.5 mL EP tube) and stored at -80°C until measurements. Samples were thawed in an ice bath and vortexed, followed by centrifugation at 3000 rpm for 5 min at 4°C. Supernatants were collected for further cytokine assays. Levels of the following 27 cytokines were analyzed using a BioPlex Pro™ Human Cytokine 27-plex Assay kit (#M500KCAF0Y, Bio-Rad, Hercules, CA, USA) in accordance with the manufacturer’s instructions: L-1β, IL-1α, IL-2, IL-4, IL-5, IL-6, IL-7, IL-8, IL-9, IL-10, IL-12(p70), IL-13, IL-15, IL-17, Eotaxin, Basic FGF, G-CSF, GM-CSF, IFN-γ, IP-10, MCP-1, MIP-1α, MIP-1β, PDGF-BB, RANTES, TNF-α and VEGF. Mean fluorescence intensities of the 192 salivary samples and 8 standards were detected via a Luminex FLEXMAP 3D System (Luminex Corp., Austin, TX, USA). Cytokine concentrations were calculated by xPONENT build 4.2.1441.0 (Luminex Corp.) using a five-parameter fit algorithm. Values obtained from the reading of samples below the sensitivity limit of detection (LOD) or above the upper limit of the sensitivity method were interpolated using a CUBIC SPINE interpolation to calculate cytokine concentrations.

### Statistical analyses

All statistical analyses were performed using R software (version 3.6.2). PCoA analysis on a range of distance metrics was performed in R using the vegan and ape package. Quantifications of variance explained in plaque microbiome, metabolome and salivary cytokines profiles were calculated using PERMANOVA with the “adonis” function in the R package vegan (as shown in **Fig. S1**). The total variance explained by each variable was calculated independently of other variables, and should thus be considered the total variance explainable by that variable. The differential abundance analyses of all measurement types were tested. First, an appropriate transformation/normalization method was applied: central-log-ratio (CLR) transformation for microbial taxonomic profiles. The transformed abundances were then used to perform differential abundance analyses between time points or groups using custom R functions (at https://github.com/shihuang047/crossRanger). To construct the co-occurrence network of molecular features from the multi-omics datasets, we identified significant associations between them using the Spearman correlation (|rho|>0.6; FDR *p*<0.05). Network was visualized in Cytoscape (Version 3.7.1). The code and all the datasets used in this study are publicly available at http://mse.ac.cn/SoH.html.

## General

We thank Jiahui Li and Duane Charbonneau for their support of this work.

## Author contributions

The project was conceptualized by J.L., J.X. and T.H.. Data collection was performed primarily by F.Y., X.L. and T.H.. Data analysis and interpretation were mainly performed by S.H., J.X., T.H., P.Z., V.X., S.W., G.J. and F.Y.. J.X., S.H., T.H., J.L., F.Y., V.X., S.W. and L.J. wrote the manuscript. All authors approved the final submission.

## Data and materials availability

All data needed to evaluate the conclusions in the paper are present in the paper and/or the Supplementary Materials. Additional data related to this paper may be requested from the authors.

