## Supplemental figures and tables for "Longitudinal multi-omics along gingivitis development reveal a suboptimal-health gum state with periodontitis-like microbiome"

### Supplementary Materials

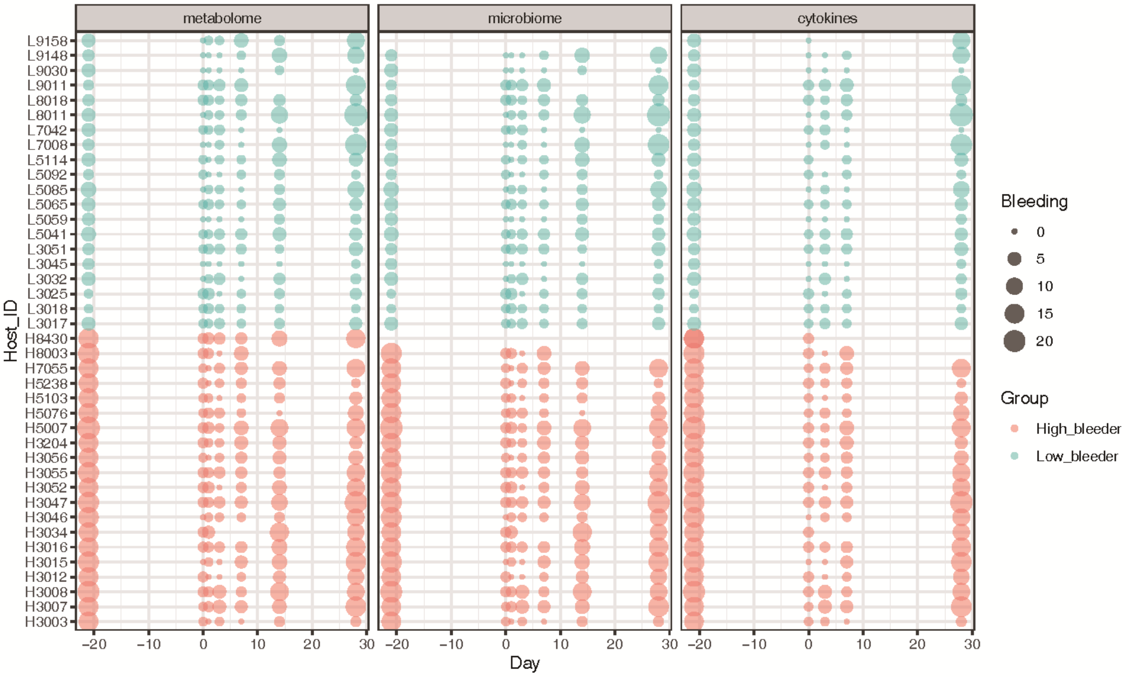


**Fig. S1. Key features of the 40-individual cohort.** Measurements (plaque metabolome, microbiome and salivary cytokines) available over time for each participant.

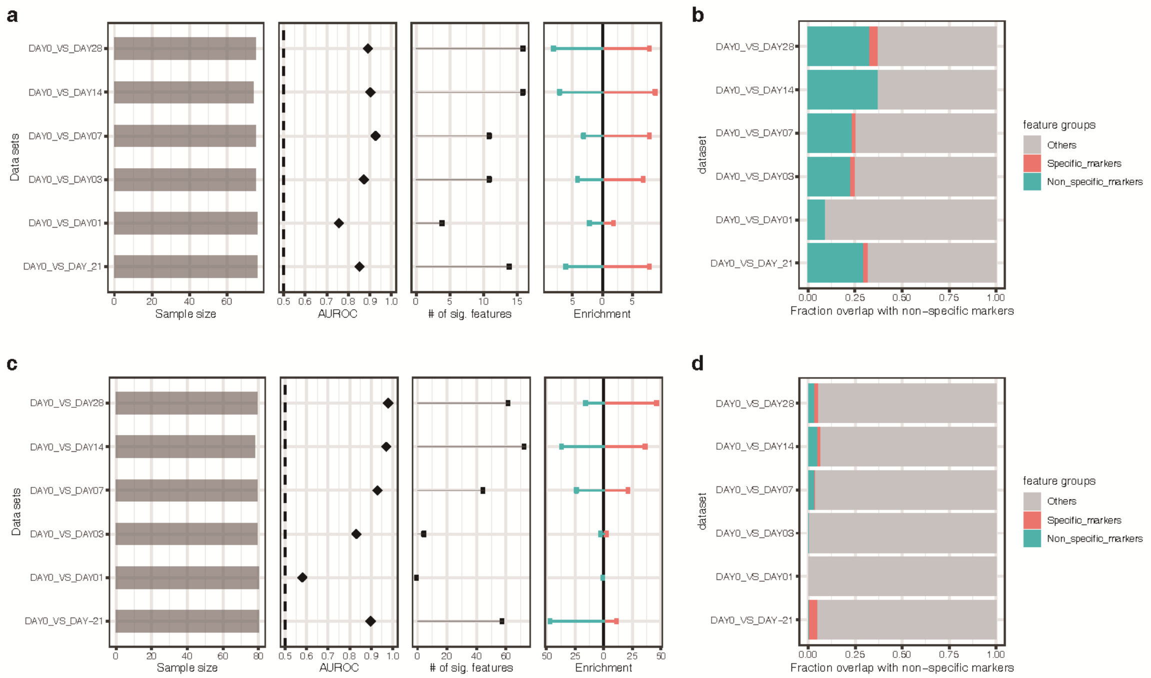


**Fig. S2**. **Temporal profiles of microbiome structure and metabolome of dental plaque unveiled the presence of a SoH stage.** (**a**) The shifts in plaque microbiota indicate a SoH stage that takes place earlier than the shifts in clinical symptoms. (**b**) The fraction of microbial taxonomical markers transiently and persistently associated with gingivitis onset and progression in this 40-adult cohort. (**c**) The dynamics of plaque metabolome also indicate a SoH stage that emerges earlier than the manifestation of clinical symptoms. (**d**) The fraction of metabolite markers transiently and persistently associated with gingivitis onset and progression in this 40-adult cohort.

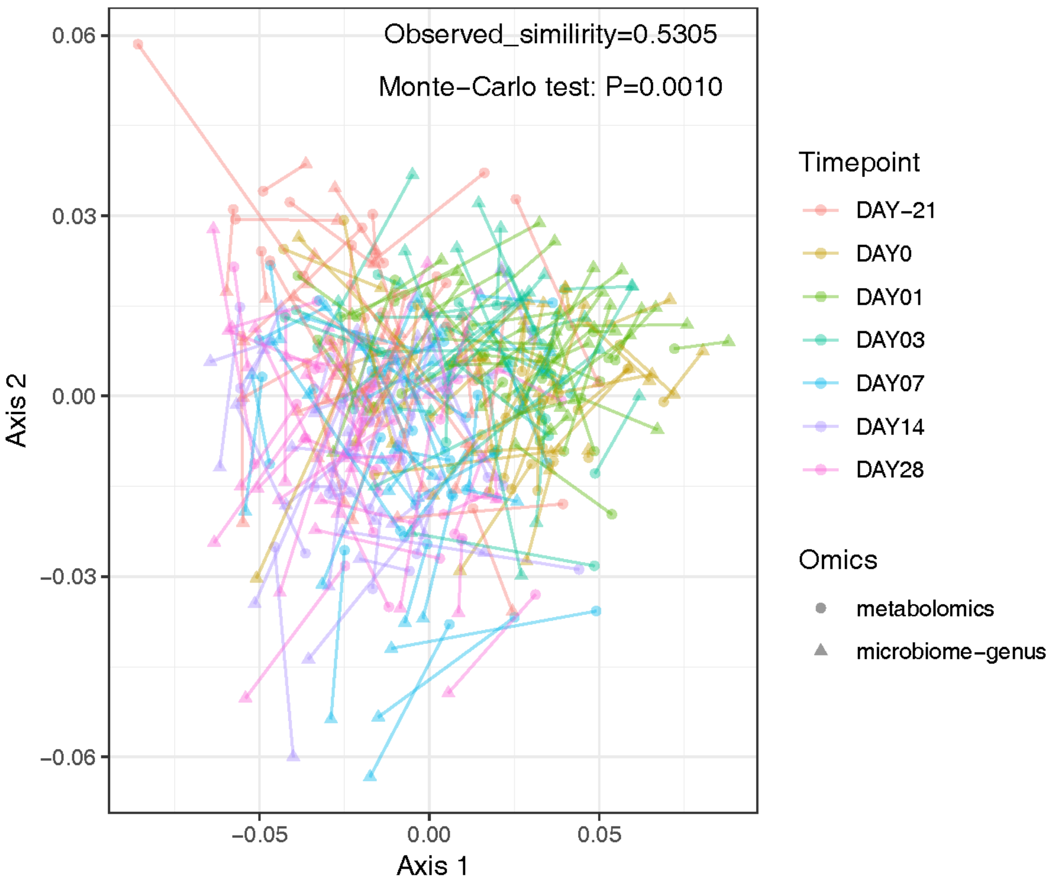


**Fig. S3. The high level of agreement between the dynamics of metabolome and microbiota along the 48-day NG-Baseline-EG process as revealed by Procrustes analysis.** In total, 261 plaque samples were simultaneously measured for metabolomics and microbiome-structure profiles. Each point represents a plaque microbiota and was colored according to the clinical status. The arrow end of each line was connected to the 16S rRNA data of a sample, whereas the other end to the corresponding plaque metabolome data. The fit of each Procrustes transformation over the first four dimensions was reported as the *p* value by 10000 Monte Carlo label permutations. A strong correlation between microbiome and metabolome was found along the 48-day process of gingivitis retrogress, onset and progression (rho=0.53).

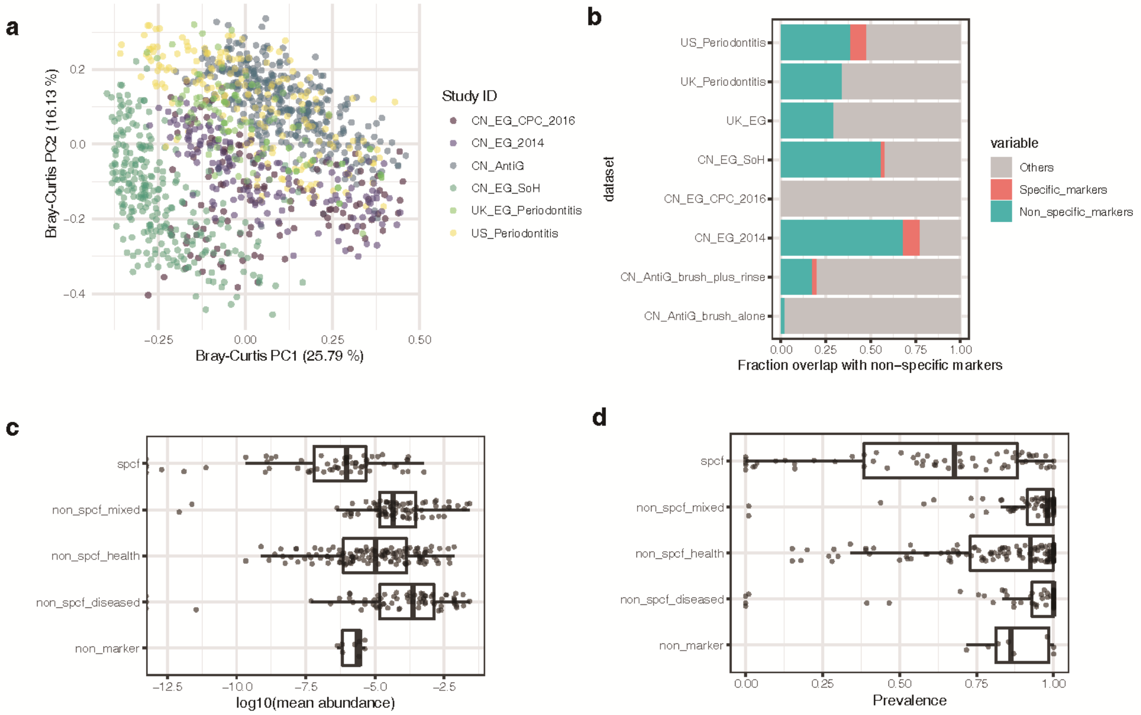


**Fig. S4. The generalizability of periodontal disease models revealed by cross-application of random forests models. (a)** PCoA of all plaque microbiomes in the meta-analysis. Each dot represents a plaque profile at the genus level and is colored per study ID. (**b**) The majority of disease-associated microbiome associations overlap with a non-specific microbial response to the periodontal disease. The bar plot indicates non-specific and disease-associated genera (green), dataset-specific disease-associated genera (red), and non-markers (grey) found in all datasets. Non-specific genera are associated with health (or disease) in at least two datasets. Dataset-specific genera are associated with health (or disease) in only one study, or associated with health (or disease) in the different direction for all the other studies. (**c and d**) The abundance and prevalence of the "non-specific genera" in all microbiomes in all data sets. Non-specific genera on the x-axis are as defined above.

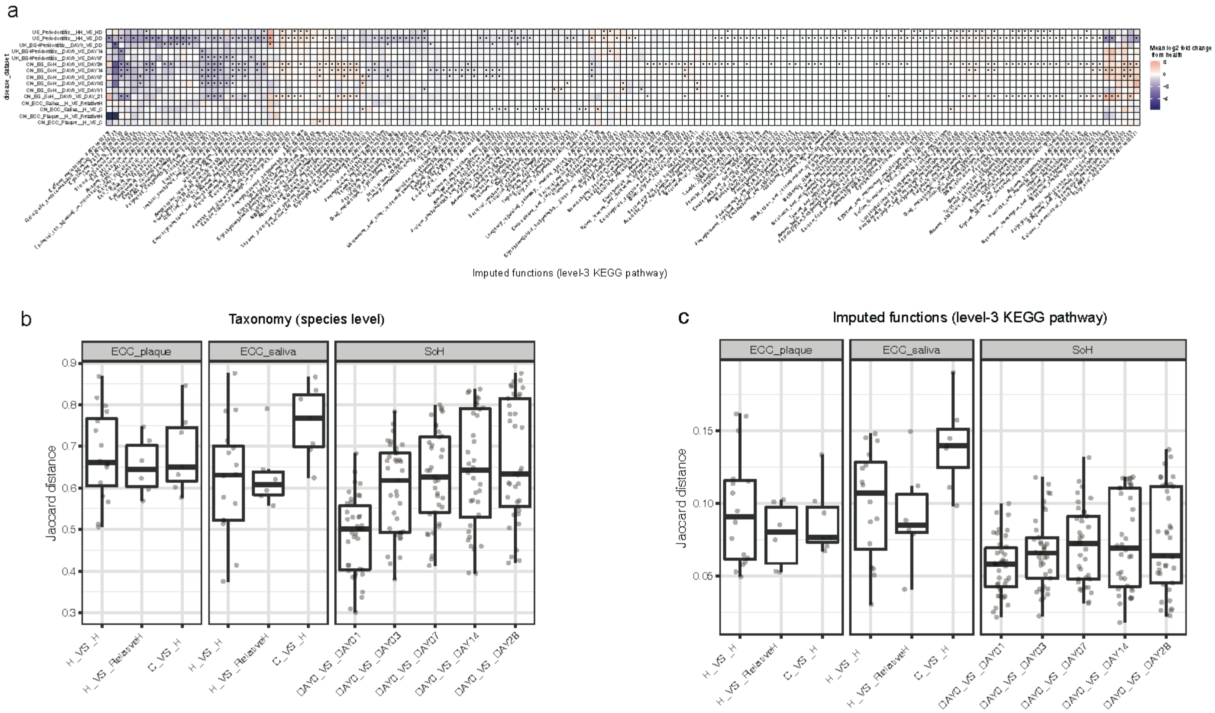


**Fig. S5. The conservation in microbiome functional change associated with the development of periodontal disease as compared to that with the development of ECC. (a)** The heatmap for the mean log2 fold changes of microbial metabolic functional pathways (with significance threshold Bonferroni-corrected $p<0.05$) in the plaque during the onset and progression of EG and ECC respectively. Asterisk: Bonferroni-corrected statistical significance (* $p \leq 0.05$). No asterisk: no significant change. The boxplots indicate the individualized distances based on taxonomic compositions (species-level; **b**) or predicted functional pathways (**c**) between each pair of sampling groups related to disease development.

**Table. S1. Clinical study design.** This is a single-leg, examiner-blind and randomized clinical study. Forty healthy adult volunteers who meet the inclusion criteria were recruited for this study. The procedures taken at each time points along 48-day NG-Baseline-EG process were highlighted via 'X' below. See **Methods** and **Fig. 1a** for additional details.

| Phase | Oral Hygiene | | | Induced Gingivitis | | | | | | Recovery |
|---|---|---|---|---|---|---|---|---|---|---|
| Procedure | Day -21 | Day -14 | Day -7 | Day 0 | Day 1 | Day 3 | Day 7 | Day 14 | Day 28 | Follow-up |
| Informed Consent | X | | | | | | | | | |
| Medical History | X | | | | | | | | | |
| Demographics | X | | | | | | | | | |
| Inclusion/Exclusion | X | | | | | | | | | |
| Continuance Criteria | | X | X | X | X | X | X | X | X | |
| saliva | X | | | X | X | X | X | X | X | X |
| Periodontal Examination | X | | | | | | | | | |
| Mazza Gingival index | X | X | X | X | x | x | X | X | X | X |
| Gingival plaque collection | X | | | X | X | X | X | X | X | |
| Dental Prophylaxis | X | X | X | | | | | | | |
| Tooth polishing | X | | X | | | | | | | X |
| Daily Supervised Oral Hygiene | X | X | X | X | X | X | X | X | | X |
| Adverse Event Recording | X | X | X | X | X | X | X | X | X | |
| Subject Accountability | | | | | | | | | X | |

**Table S2. Overview of the 40-individual cohort that underwent the controlled oral hygiene regimen.** (**a**) Overview of the cohort. (**b**) Background information on each subject that participated in the cohort.

| Demographic/Statistic Category | High_bleeder (n=20) | Low bleeder (n=20) | Overall (n=40) | *p*-value |
|---|---|---|---|---|
| **Age (Years)** | | | | |
| Mean (SD) | 33.60 (6.52) | 37.65 (9.44) | 35.63 (8.27) | 0.123[a] |
| Min.-Max. | 23 - 52 | 22 - 53 | 22 - 53 | |
| **Gender** | | | | |
| Female[b] | 16 (80%) | 16 (80%) | 32 (80%) | 1.000[c] |
| Male[b] | 4 (20%) | 4 (20%) | 8 (20%) | |
| **Smoking** | | | | |
| Yes[b] | 2 (10%) | 2 (10%) | 4 (10%) | 1.000[c] |
| No[b] | 18 (90%) | 18 (90%) | 36 (90%) | |

[a] Two-sided ANOVA *p*-value for the treatment comparison.

[b] The number (percent) of subjects in each category.

[c] Two-sided chi-square *p*-value for the treatment comparison.

[d] Two-sided Fisher's exact test *p*-value for the treatment comparison.

| Host_ID | Group | Age | Gender | Smoking |
|---|---|---|---|---|
| 3003 | High_bleeder | 38 | F | N |
| 3007 | High_bleeder | 37 | F | N |
| 3008 | High_bleeder | 29 | F | N |
| 3012 | High_bleeder | 30 | F | N |
| 3015 | High_bleeder | 33 | F | N |
| 3016 | High_bleeder | 30 | F | N |
| 3017 | Low_bleeder | 28 | F | N |
| 3018 | Low_bleeder | 22 | F | N |
| 3025 | Low_bleeder | 41 | F | N |
| 3032 | Low_bleeder | 53 | F | N |
| 3034 | High_bleeder | 52 | F | N |
| 3045 | Low_bleeder | 46 | M | N |
| 3046 | High_bleeder | 31 | M | Y |
| 3047 | High_bleeder | 31 | F | N |
| 3051 | Low_bleeder | 36 | F | N |
| 3052 | High_bleeder | 37 | M | N |
| 3055 | High_bleeder | 34 | M | N |

| | | | | |
|---|---|---|---|---|
| 3056 | High_bleeder | 28 | F | N |
| 3204 | High_bleeder | 23 | F | N |
| 5007 | High_bleeder | 36 | F | N |
| 5041 | Low_bleeder | 41 | F | N |
| 5059 | Low_bleeder | 47 | F | N |
| 5065 | Low_bleeder | 31 | F | N |
| 5076 | High_bleeder | 33 | F | N |
| 5085 | Low_bleeder | 49 | F | N |
| 5092 | Low_bleeder | 31 | M | Y |
| 5103 | High_bleeder | 31 | F | N |
| 5114 | Low_bleeder | 31 | F | N |
| 5238 | High_bleeder | 47 | F | N |
| 7008 | Low_bleeder | 46 | F | N |
| 7042 | Low_bleeder | 47 | F | N |
| 7055 | High_bleeder | 30 | F | N |
| 8003 | High_bleeder | 30 | M | Y |
| 8011 | Low_bleeder | 30 | F | N |
| 8018 | Low_bleeder | 23 | M | N |
| 8430 | High_bleeder | 32 | F | N |
| 9011 | Low_bleeder | 29 | M | Y |
| 9030 | Low_bleeder | 31 | F | N |
| 9148 | Low_bleeder | 47 | F | N |
| 9158 | Low_bleeder | 44 | F | N |
